# Electrophysiological properties of mesodiencephalic junction neurons projecting to the inferior olive

**DOI:** 10.64898/2026.04.10.717667

**Authors:** S. Voerman, X. Wang, L.W.J. Bosman, R. Broersen, C.I. De Zeeuw

## Abstract

Neocortex has the largest volume, while cerebellar cortex houses most neurons of the mammalian brain, underscoring paramount functions for interactions between them. Neocortex projects to the cerebellum via both the mossy fibre and climbing fibre system; whereas the mossy fibres find their origin in the pons, the climbing fibres are relayed via the subnuclei of the inferior olive (IO) that receive their cortical input from the mesodiencephalic junction (MDJ). Since climbing fibres regulate cerebellar plasticity in a timing-dependent manner, the MDJ-IO pathway probably contributes to both sensorimotor and cognitive learning. However, the physiological properties of IO-projecting MDJ neurons remain largely unknown. Here, we made targeted whole-cell recordings in acute brain slices of the murine MDJ, separating IO- from non-IO-projecting neurons following retrograde tracing. We show that IO-projecting neurons are spontaneously active and are capable of high frequency-bursts up to 350 Hz during current injections. Action potentials of IO-projecting MDJ neurons depolarize and re-polarize quickly, and hyperpolarizing inputs consistently trigger rebound action potentials upon stimulus offset. Instead, non-IO-projecting neurons in the same MDJ region are hardly spontaneously active with little rebound activation. Even so, IO-projecting MDJ neurons can also transform excitatory and inhibitory inputs into a proportional output, indicating that they can also employ rate coding. Moreover, neocortical inputs to IO-projecting MDJ neurons can directly be integrated with their inputs from the cerebellar nuclei. Our results highlight how neocortical inputs to the MDJ can be transformed into diverse IO firing patterns and thereby contribute to the required specifics of cerebellar learning.

**Key points summary:**

- The timing of complex spikes in cerebellar Purkinje cells is essential for cerebellar motor learning and proper timing is achieved through the modulation of activity in the inferior olive, the source of all climbing fibres.
- A major source of excitatory input to the inferior olive is the mesodiencephalic junction.
- We find that that mesodiencephalic junction neurons projecting to the inferior olive represent a population of physiologically distinct neurons, capable of high firing frequencies and rebound activity following inhibition.
- We reveal how mesodiencephalic junction neurons process excitatory inputs from the neocortex and cerebellar nuclei, as well as inhibitory inputs, enabling mesodiencephalic junction neurons to bidirectionally regulate their activity.
- These findings shed light on how the inferior olive, critical for cerebellar motor learning, is controlled by the mesodiencephalic junction.

## Introduction

The cerebellum has long been known to be vital for motor learning and is increasingly also recognized as a key player in non-motor forms of learning (Raymond et al., 1996; Timmann et al., 2010; De Zeeuw & Ten Brinke, 2015; Hull, 2020). To facilitate its role in learning, the cerebellum receives inputs from many different regions of the cerebral cortex, including sensory, motor and associative areas (Suzuki et al., 2012; Keser et al., 2015; Palesi et al., 2017; Wang et al., 2022). Like all inputs to the cerebellar cortex, neocortical information reaches the cerebellum mostly via two routes, involving either the mossy fibres or the climbing fibres. Neocortical signals to the mossy fibres are relayed primarily via the pons, and those to the climbing fibres via the inferior olive (IO). As direct projections from the neocortex to the IO are relatively sparse, it is likely that most of the information passes through an intermediate station (Swenson et al., 1989; Ruigrok et al., 2015). Given its anatomical pathways, the mesodiencephalic junction (MDJ) could play this role as a bridge between the forebrain and IO, and thus channels neocortical input to the climbing fibres that target the cerebellum (Sasaki et al., 1975; Apps & Watson, 2013; Ruigrok et al., 2015; Wang et al., 2022; Broersen & De Zeeuw, 2024). When active, the olivary climbing fibres evoke complex spikes in the Purkinje cells of the cerebellar cortex (Eccles et al., 1966; De Zeeuw et al., 2011). Well-timed complex spike firing is critical for the regulation of synaptic plasticity in Purkinje cells, and thereby also for cerebellar learning (Ito et al., 1982; Coesmans et al., 2004; Gao et al., 2012; De Zeeuw & Ten Brinke, 2015; Suvrathan et al., 2016; Bouvier et al., 2018; Romano et al., 2018). Hence, to understand cerebellar learning, we have to understand the mechanisms that drive activity in the IO.

The neuropil of the IO is almost entirely devoid of interneurons, but its subnuclei are organized in clusters of neurons that are strongly connected by dendrodendritic gap junctions located inside isolated glomeruli with strategically located excitatory and inhibitory inputs (Llinas et al., 1974; Sotelo et al., 1974; De Zeeuw et al., 1989; Fredette & Mugnaini, 1991; Loyola et al., 2019; Vrieler et al., 2019; Negrello et al., 2019). In addition to this unique anatomical arrangement, neurons in the IO also have distinct electrophysiological characteristics. Intrinsic Ca2+ and K+ conductances can generate sub-threshold oscillations of the membrane potential in large portions of the IO neurons (Khosrovani et al., 2007; Choi et al., 2010; Negrello et al., 2019; Loyola et al., 2023). During the depolarized phase of the subthreshold oscillation, the chance of action potential firing is increased, but in the awake brain this effect can be mitigated by synaptic input (Negrello et al., 2019; Loyola et al., 2023). The abundant gap junctions affect the properties of the subthreshold oscillations and promote synchrony between IO neurons (Llinas et al., 1974; Bazzigaluppi et al., 2012; Negrello et al., 2019). The latter is important for the reliability of cerebellar encoding, and abolition of coupling and synchrony can lead to motor deficits (Van Der Giessen et al., 2008; De Gruijl et al., 2014; Tsutsumi et al., 2019; Ikezoe et al., 2023, 2023).

In the awake brain, synaptic inputs have a strong impact on the intrinsic dynamics of IO neurons (Lang et al., 1996; Lang, 2002; Best & Regehr, 2009; Negrello et al., 2019; Loyola et al., 2023; Guo & Uusisaari, 2025). The major source of GABAergic input is the nucleo-olivary pathway, connecting specific neurons in the cerebellar nuclei with the IO (De Zeeuw et al., 1989, 1998). Glutamatergic input to large parts of the IO, i.e., that of the principal olive and rostral medial accessory olive, primarily comes from the MDJ (Fig 1a) (Onodera, 1984; De Zeeuw et al., 1989, 1998; Ruigrok et al., 2015; Wang et al., 2022). The MDJ itself receives extensive inputs from a widely varying range of brain regions, including the primary and secondary somatosensory cortex, the primary and secondary motor cortex, the basal ganglia, ascending sensory regions, and the cerebellar nuclei (Kubo et al., 2018; Wang et al., 2022; Ruigrok et al., 2023). Thus, the MDJ is well-positioned to be the main intermediate between the forebrain and the IO, which makes it well positioned to be an essential component of the cerebro-cerebellar learning network.

**Figure 1.**
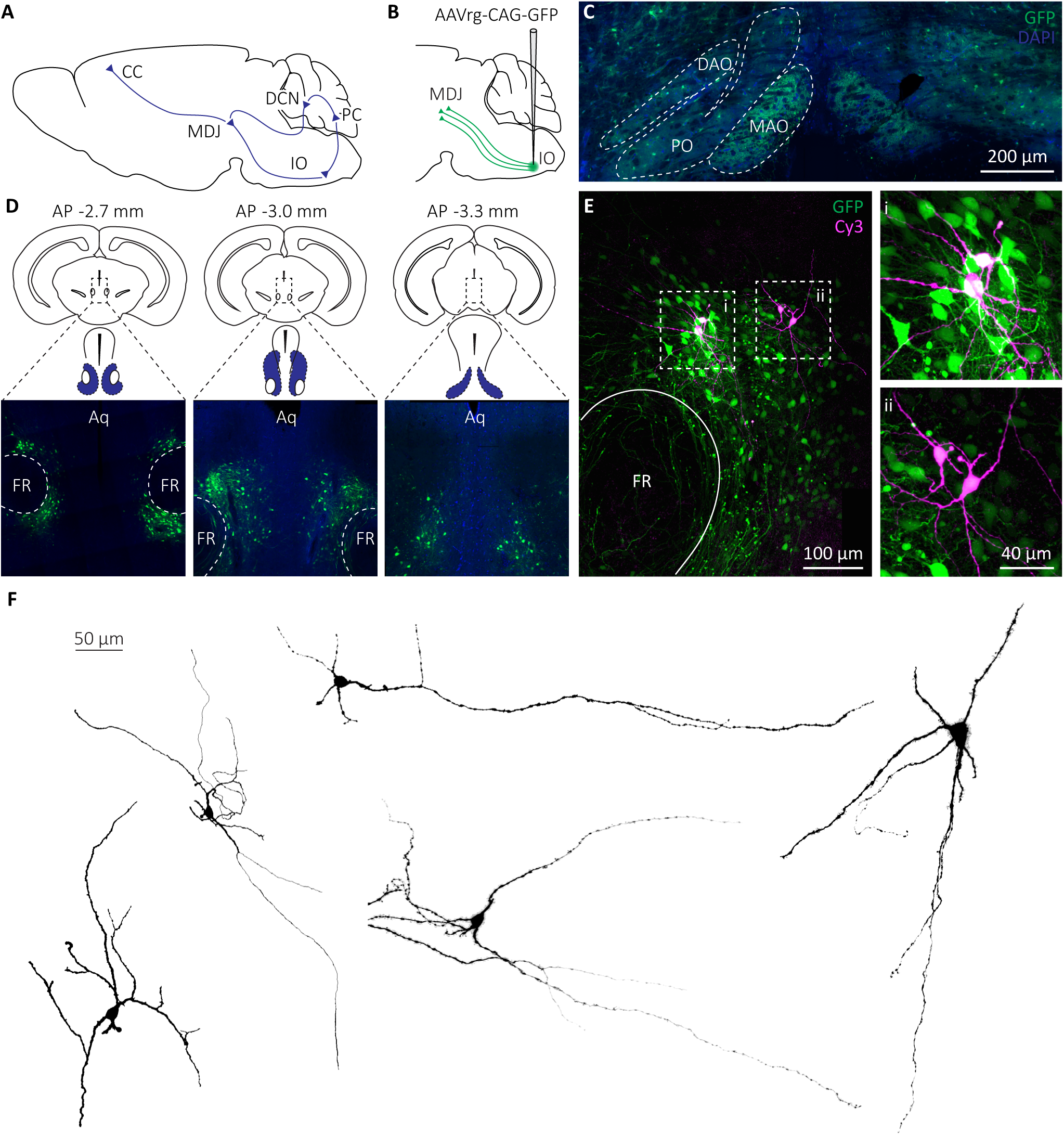
Overview of experimental design. **(A)** Scheme of the connectivity of the MDJ. The MDJ receives input from the cerebral cortex (CC) and the deep cerebellar nuclei (DCN), and projects to the inferior olive (IO). **(B)** The injections in the IO with AAVrg-CAG-GFP. Injections were bilateral (not shown). (C) Confocal image of the injections in the IO, showing the expression in the dorsal accessory olive (DAO), medial accessory olive (MAO), and the principal olive (PO). **(D)** Confocal images with examples of GFP expression after AAVrg injection in the IO, at three different anterior-posterior (AP) positions. Expression centres around the fasciculus retroflexus (FR) and below the aqueduct (Aq). **(E)** Confocal image made upon completion of whole-cell recordings, confirming the location of the recorded neurons in the MDJ. Expression of GFP again centres around the FR. Recorded neurons have been enlarged in panels i and ii. **(F)** Morphological reconstructions of 5 example IO-projecting MDJ neurons. Scale is the same for all.

Despite its potential importance for understanding brain-wide learning processes, the physiology of MDJ neurons remains unknown. Therefore, we investigated the anatomical and electrophysiological properties of MDJ neurons, comparing neurons that do and do not project to the IO. We found that IO-projecting MDJ neurons represent a relatively homogenous neuronal population with physiological properties distinct from their non-IO-projecting neighbours. IO-projecting neurons were spontaneously active, exhibited rebound spikes after cessation of strong inhibitory input, and were capable of high firing rates, in contrast to most non-IO projecting neurons. Optogenetic stimulation experiments further demonstrated that individual IO-projecting neurons can be excited by different regions of the neocortex, as well as the deep cerebellar neurons, indicating that the MDJ serves as a critical hub between the neocortex and olivo-cerebellar system.

## Methods

### Animals

Both male and female C57BL/6J mice were used (41 mice, M = 22, F = 19). For the visualization of inhibitory synaptic markers, we used a GAD2-Cre-Ai14 mouseline (2 mice, M=1, F=1). All mice used in experiments were older than P50 (P88-221, mean = P122). All animal experiments were performed after obtaining permission from the Dutch national authority on animal experiments (Centrale Commissie Dierproeven, The Hague, The Netherlands, license no. AVD1010020 2417861), in accordance with Dutch and EU regulations and overseen by the institutional Animal Welfare Board of Erasmus MC.

### Viral injections

All surgeries were performed under isoflurane anaesthesia, with an induction concentration of 2%– 4% and a maintenance concentration of 1%–2% (v/v) in oxygen, at a flow rate of 0.8 L/min. Carprofen (Rimadyl, Pfizer) and buprenorphine (Temgesic, Indivior, Richmond, VA, USA) were subcutaneously administered at 10 µL/g body weight 15 minutes prior to surgery. Sterile eye ointment was applied to prevent corneal drying. Body temperature was maintained and monitor using a homeothermic temperature controller. Injections were conducted using Hirschmann® microcapillary pipettes (10 µm inner diameter; Sigma-Aldrich, Germany) coupled with a Nanoject II Auto-Nanoliter Injector (Drummond Scientific Company, USA) while the mouse was placed in a Model 1900 Sterotaxic Alignment System (Kopf Instruments, USA). The hair over the designated area was carefully removed using a surgical blade. A small circular craniotomy (1 to 2 mm diameter) was performed at the target coordinates using a dental drill. For bilateral injections into the inferior olive (IO), 60 nL of retrograde AAV (pAAV-CAG-GFP, 37825-AAVrg, Addgene) was administered into the left rostral IO (anteroposterior [AP]:-2.5 mm, mediolateral [ML]: 0.3 mm, depth: 5.4 mm, relative to lambda) and another 60 nL of pAAV-CAG-GFP into the right rostral IO (AP:-2.5 mm, ML: 0.3 mm, depth: 5.4 mm, relative to lambda). For optogenetics experiments, 50 nL of pAAV-CAG-GFP (37825-AAVrg, Addgene) was injected into the rostral IO (AP: -2.5 mm, ML: 0.3 mm, depth: 5.4 mm, relative to lambda). Additionally, 100 nL of pAAV-Syn-Chronos-GFP (59170-AAVG5, Addgene) was delivered into the deep cerebellar nuclei (DCN), including the interposed nucleus (IN) and dentate nucleus (DN) (AP: -2.2 mm, ML: -2.3 mm, depth: 2.4 mm, relative to lambda). A volume of 150 nL of pAAV-Syn-ChrimsonR-tdT (591171-AAV9, Addgene) was injected into the primary/secondary motor cortex (M1/M2) (AP: 1.2 mm, ML: 1.0 mm, depth: 0.6 mm, relative to bregma) and another 150 nL into the whisker primary somatosensory cortex (wS1) (AP:-1.5 mm, ML:-3.5 mm, depth: 0.6 mm, relative to bregma). Following viral injection, Kwik-Sil (World Precision Instruments [WPI], USA) was applied to seal the craniotomy site. Half-dry dental cement (Super-Bond C&B, Sun Medical, Japan) was then applied over the skull for stabilization. Postsurgical pain was managed with carprofen (Rimadyl®, Pfizer) and buprenorphine (Temgesic, Indivior, Richmond, VA, USA) for two days during the postoperative recovery period.

### In vitro slice electrophysiology

After a survival time of at least 3 weeks following virus injection, mice were decapitated under isoflurane anaesthesia, and their brains were removed and moved into an ice-cold slicing solution containing (in mM): 240 sucrose, 2.5 KCl, 1.25 Na2HPO4, 2 MgSO4, 1 CaCl2, 26 NaHCO3, and 10 D-glucose. Coronal slices with a thickness of 200 µm were obtained from the midbrain area containing the MDJ using a vibratome (Leica VT1200S, Leica Biosystems). Slices were incubated in carbogenated artificial cerebrospinal fluid (ACSF) containing in mM: 124 NaCl, 2.5 KCl, 1.25 Na2HPO4, 2 MgSO4, 2 CaCl2, 26 NaHCO3, and 15 D-glucose for 1 hour at ±34 °C, and kept for a further 6-8 hours in ACSF at ± 21 °C. After the incubation period, slices were moved to an electrophysiology rig for ex vivo whole cell recordings. All electrophysiology experiments were performed with an EPSC10-USB amplifier (HEKA electronics, Germany) using Patchmaster software (version 92x2). Acquired signals were sampled at 50 kHz. Slices were visualized with an upright microscope (Zeiss AXIO imager M2, Carl Zeiss Microscopy, Göttingen, Germany). LED triggers for fluorescence visualization and optogenetic stimulation were performed with a Colibri 7 system (Carl Zeiss Microscopy, Gottingen, Germany).

Recording pipettes with a resistance between 2-5 MΩ were used (OD 1.65 mm, ID 1.11 mm, World Precision Instruments, Sarasota, FL, USA). The pipettes were prepared using a P-1000 micropipette puller (Sutter instruments, Novato, CA, USA). For all experiments except for sIPSC recordings, pipettes were filled with a K-gluconate internal solution containing (in mM): 120 K-gluconate, 9 KCl, 10 KOH, 4 NaCl, 3.48 MgCl2, 10 HEPES, 28.5 Sucrose, 4 Na2ATP, 0.4 Na3GTP (pH 7.25-7.35, osmolarity 300 ± 5 mOsm). The internal solution also contained biocytin (Sigma Aldrich) at a concentration of 1 mg/ mL, which was used for the post-hoc visualization of recorded neurons (see immunohistochemistry for details). Recordings of sIPSCs were done using a caesium-chloride internal containing in mM: 150 CsCl, 1.5 MgCl2, 0.5 EGTA, 10 HEPES, 4 Na2ATP, 0.3 Na3GTP, 5 Qx-314. sIPSCs were recorded with 1 µM TTX (HelloBio, Bristol, United Kingdom), 20 µM NBQX (Tocris Bioscience, Bristol, United Kingdom), and 50 µM D-AP5 or DL-AP5 (Tocris Bioscience).

The ratio between series and input resistance was generally lower than 20%, and series resistance was mostly below 30 MΩ. Holding current required to maintain voltage clamp differed substantially per cell, but was roughly between 0 and -250 pA. Bridge balance and pipette capacitance compensation was used during all current-clamp experiments. Upon completion of the electrophysiological recordings, slices were removed from the setup and kept in 4% paraformaldehyde (PFA) at 4 °C for at least 24 hours for subsequent histological analysis.

### Experimental design

#### Electrophysiological characterization

In the brain slices, either GFP-positive or-negative MDJ neurons were targeted for electrophysiological recording. Cells expressing GFP were considered as projecting from the MDJ to the IO, whereas cells that did not express GFP within the same region, were considered as putative non-IO-projecting neurons. Once whole-cell configuration was achieved, cells were voltage-clamped at-65 mV. We recorded the following protocols for every cell: resistance test, spontaneous PSCs, current injections, slow ramp current injections, and spontaneous firing. Series and input resistance was determined with a voltage step of-10 mV for 100 ms, again from -65 mV. Membrane capacitance was also calculated using the same -10 mV voltage step. The decay curve of the -75 to -65 mV step was fitted to a double exponential function, from which the time constant (τ) was determined. The longest τ was used to calculate membrane capacitance.

Current injections were adapted to the input resistance of the neuron. Neurons with low (< ∼250 MΩ), medium (∼250-500 MΩ), or high (> ∼500 MΩ) input resistances were injected with-500 to 500, -250 to 250, or-100 to 100 pA, respectively. Current injections were made with steps of 10 pA. The starting current was set by the amount of current to keep membrane voltage at ∼-65 mV.

Slow ramping currents consisted of a ramping current injection, increasing by 1 pA/s. This was continued until at least 10 action potentials were evoked, which were used in the analysis of spike properties (i.e. spike threshold). Spontaneous firing behaviour was recorded by setting the injected current in current-clamp at 0 pA for 20 s. Cells that fired very few action potentials (frequency < 0.1 Hz) were not considered as spontaneously active.

#### Spontaneous IPSCs

Spontaneous IPSCs were recorded at a membrane potential of-65 mV. NBQX disodium salt (10-20 µM) and DL-AP5 (50 µM) were added to the extracellular solution to block AMPA and NMDA receptors, respectively. In some experiments, picrotoxin (100 µM) (Sigma-Aldrich) was washed in to determine if spontaneous IPSCs were GABA-mediated. Wash-in recordings consisted of a baseline measurement of 2 min, followed by up to 20 min of additional recordings after the start of wash-in.

#### Optogenetics

Mice transfected with ChrimsonR and Chronos were sacrificed, and brain slices were obtained as described earlier. Brain slices were protected from ambient light during incubation and recording. ChrimsonR and Chronos were activated using LED stimulation (Zeiss Colibri 7) with a wavelength of 631±33 nm or 469±38 nm, respectively (Klapoetke et al., 2014). All optogenetic experiments were performed in the presence of 1 µM TTX and 200 µM 4-AP (Tocris). To measure the impact of activation of both ChrimsonR and Chronos on single MDJ neurons, we used a dichroic mirror (DLP 710). No additional excitation or emission filters were used. Blue and red light stimuli were given within 500-1000 ms intervals, at a frequency of 1 Hz for 10 ms each. If a double response to both blue and red light stimulation was observed, an initial 500 ms red light stimulus was given to deplete the synapses of ChrimsonR expressing fibres. After a recovery time of 1.5s a blue light stimulus was given to test any remaining Chronos response.

### Immunohistochemistry

Previously recorded neurons that were filled with biocytin during electrophysiological experiments were kept in 4% paraformaldehyde (PFA) for at least 24 hours at 4 °C. Brain slices were washed 3 times for 10 min in PBS. Slices were then kept in a solution containing 1:400 streptavidin Cy3 or Cy5 (Jackson ImmunoResearch, Camebridgeshire, United Kingdom, RRIDs: AB_2337244 and AB_2337245), 2% NHS, and 0.4% triton in PB for at least 4 hours at room temperature. Subsequently, slices were washed with PB (0.05M) 3 times for 10 minutes, stained with DAPI (ThermoFisher Scientific, Cat. No. D3571), and washed another 3 times with PB. Slices were mounted on microscope slides, and covered with cover slips for imaging. The quality of viral injections was confirmed by inspecting 50 µm thick slices of gelatine-embedded brain regions that were stained with DAPI.

For the visualization of the inhibitory synaptic markers VGAT and gephyrin, two GAD2-Cre-Ai14 mice (CAG-tdTomato-14 x GAD2-Cre on B6 background) were perfused with 4% PFA, and 50 µm thick coronal sections were cut on a cryostat (Leica SM2000R, Leica Biosystems). Slices were washed in phosphate buffer, sodium citrate, and once more in phosphate buffer for 10 minutes each. Slices were then blocked in a blocking solution containing 10% NHS, and 0.5% triton for 60 minutes. The primary antibodies used were VGAT (1:1000, guinea pig, Synaptic Systems, Göttingen, Germany, Cat. No. 131004, RRID: AB_887886) and gephyrin (1:2000, mouse, Synaptic Systems, Cat. No. 147011, RRID: AB_887717). Slices were incubated in primary antibodies for 24h at 4 °C. After incubation, slices were washed with phosphate buffered saline three times for 10 minutes. Secondary antibodies used were mouse Cy5 (1:400, Jackson ImmunoResearch, Cambridge, United Kingdom) and guinea pig Alexa-Fluor 488 (1:400, Jackson ImmunoResearch, RRID: AB_2337245). Slices were incubated in secondary antibodies with 2% NHS and 0.4% triton in phosphate buffered saline for at least 2 hours at ±21 °C. Slices were again washed with PB, and stained with DAPI, followed by a wash with PB, each for 10 minutes. Finally, slices were microscope slides, and covered with cover slips for imaging.

### Microscopy and imaging

Brain slices were imaged with an upright confocal (Zeiss LSM700, Carl Zeiss Microscopy, Gottingen, Germany) or fluorescence microscope (Zeiss Axioscope). Cy3/5 labelled neurons were visualized to determine the location and to confirm their expression of GFP. Excitation was at 405, 488, 555 and 639 nm. Fluorescent microscope images were taken at a magnification of 10x or 20x. Z-stack confocal images used for neuronal tracings were taken using a magnification of 40x or 63x with a z step size of 0.75 – 2.0 µm. Neuronal tracing was performed with Fiji/ImageJ using the SNT plugin (https://imagej.net/plugins/snt/). VGAT/Gephyrin images were taken at a magnification of 100x.

### Data exclusion

Electrophysiological data used for the electrophysiological characterization of GFP+ and GFP- MDJ neurons were excluded when one or more of the following criteria were met. First, if the injection site aimed at the IO was not clearly hitting any olivary subnucleus, cells from that mouse were excluded from analysis. Second, cells were excluded if the access resistance (Ra) was greater than 30 MΩ or the ratio between the access resistance and the input resistance (Ri) was greater than 0.2 (Ra > 20% of Ri). These values indicate a bad recording quality. One neuron in the GFP-negative group with an input resistance of ∼2500 MΩ was excluded, as its input resistance exceeded all other neurons by a factor of ∼3. Finally, cells were also excluded from further analysis if the amplitude of action potentials was less than 80 mV. The access resistance, which is a measure for the technical quality of the recordings, did not differ between GFP+ and GFP- neurons (GFP+: 21.7 ± 3.80 MΩ, n = 44; GFP-: 21.2 ± 4.90 MΩ, n = 27; p = 0.651, Independent t-test). Electrophysiological data for optogenetics and sIPSC experiments were not excluded based on a cutoff of Ra or Ri.

### Data analysis

Electrophysiological data were analysed using custom Python scripts (https://www.python.org/) (Python 3.13.5). All codes used for data analyses, figure generation, or otherwise, will be made freely available upon acceptance of this manuscript on GitHub (https://github.com/s-voerman). sIPSCs were analysed automatically using a deconvolution based method (Pernía-Andrade et al., 2012). Principal component analysis was performed using Python (scikit-learn), using the following electrophysiological properties: Membrane resistance, membrane capacitance, resting membrane voltage, action potential afterhyperpolarization, action potential threshold, action potential amplitude, action potential half width, spontaneous action potential frequency, sag ratio, average rebound action potential count and average firing rate. These comprised all the electrophysiological data for which we could obtain single continuous numerical values.

### Statistical analysis

Statistical analysis was performed using Python, with the exception of linear mixed models, which were performed with R (https://www.r-project.org/) (R 4.5.2) using the lme4 and lmerTest packages. Statistical significance of fixed effects was assessed using type III ANOVA. Data were tested for equality of variance, normality, and skewness. Equality of variance was assessed using Levene’s test, and skewness was calculated by calculating the kurtosis. Skewness was used to determine the normality of distributions. If data were independent, of equal variance, and normally distributed, we used an independent samples t-test to assess differences between populations. If data were not normally distributed, we used a Mann-Whitney U test. If data were not of equal variance we used Welch’s t-test. If data were normally distributed, statistics were displayed as mean ± standard deviation (SD), and if not as median (inter-quartile range). For comparison of contingency tables we used Fisher’s exact test. Where possible, p-values are given in exact values.

The statistical analysis of data obtained from positive and negative current injections were performed using linear mixed models. This was necessary, because not all data for all current injections were available for every neuron. sIPSC wash-in recordings were also analysed with a linear mixed model, as not all neurons survived the full 20 min recording. The dependent variables were either firing rate, spike latency, mAHP, spike count, sag ratio, instantaneous firing rate, or action potential amplitude.

## Data availability

All raw and processed data produced in this article will be made available upon reasonable request to the corresponding author.

## Code availability

All codes used in the analyses of data and the generation of figures, as well as the data needed to generate the plots and figures will be made freely available on GitHub after acceptance of this manuscript (https://github.com/s-voerman). For questions, please e-mail Stijn Voerman.

## Results

### Labelling of MDJ-IO neurons

To characterize the physiological properties of IO-projecting neurons in the MDJ, we performed whole-cell recordings in acute brain slices made from mice after bilateral injection of a retrograde adeno-associated virus (AAV) in the IO (Fig 1b,c). If a neuron in the MDJ expressed GFP (GFP+ neurons), this implied that that neuron took up the virus and could, therefore, be classified as an IO-projecting neuron. Neurons that did not express GFP (GFP- neurons) were presumably targeting other areas of the brain (Fig 1d,e). However, we cannot exclude that the population of GFP- neurons also included a minority of IO-projecting neurons that did not effectively take up the injected virus or that the population of GFP+ neurons included a minority of non-IO projecting neurons that took up the virus in regions that were not the IO. We also examined GFP expression in the region surrounding the FR and observed that IO-projecting MDJ neurons had varicosities around the FR. This suggests that these neurons, in addition to their projection to the IO, may also have local projections (Fig 1e). As our recorded neurons were filled with biocytin, it was possible to reconstruct the morphology of recorded neurons (Fig 1f).

### Electrophysiological properties of MDJ-IO neurons

Next, we studied the passive membrane properties of MDJ neurons. Whole-cell voltage-clamp recordings revealed that IO-projecting (GFP+) neurons have significantly lower membrane resistances than putative non-IO projecting (GFP-) neurons in the MDJ (Fig 2a) (GFP+: 202.1 ± 78.2 MΩ, n = 44; GFP-: 439.7 ± 193.9 MΩ, n = 27; p = 1.34e-6, Welch’s t-test). We noted that the variation in membrane resistance was also lower in GFP+ than in GFP- neurons (p = 5.50e-6, Levene’s test), suggesting that MDJ-IO neurons represent a more homogeneous population than GFP- neurons in the MDJ. Other passive electrophysiological properties were similar between the GFP+ and GFP- neurons (Fig 2b,c). This was true for the membrane capacitance (Cm) (Fig 2b) (GFP+: 54.0 ± 20.6 pF, n = 44; GFP-: 49.0 ± 22.5 pF, n = 27; p = 0.344, Independent t-test) and the resting membrane voltage (Vm) (Fig 2c) (GFP+: -55.4 ± 3.24 mV, n = 44; GFP-:-59.4 ± 8.67 mV, n = 27; p = 0.057, Welch’s t-test).

**Figure 2.**
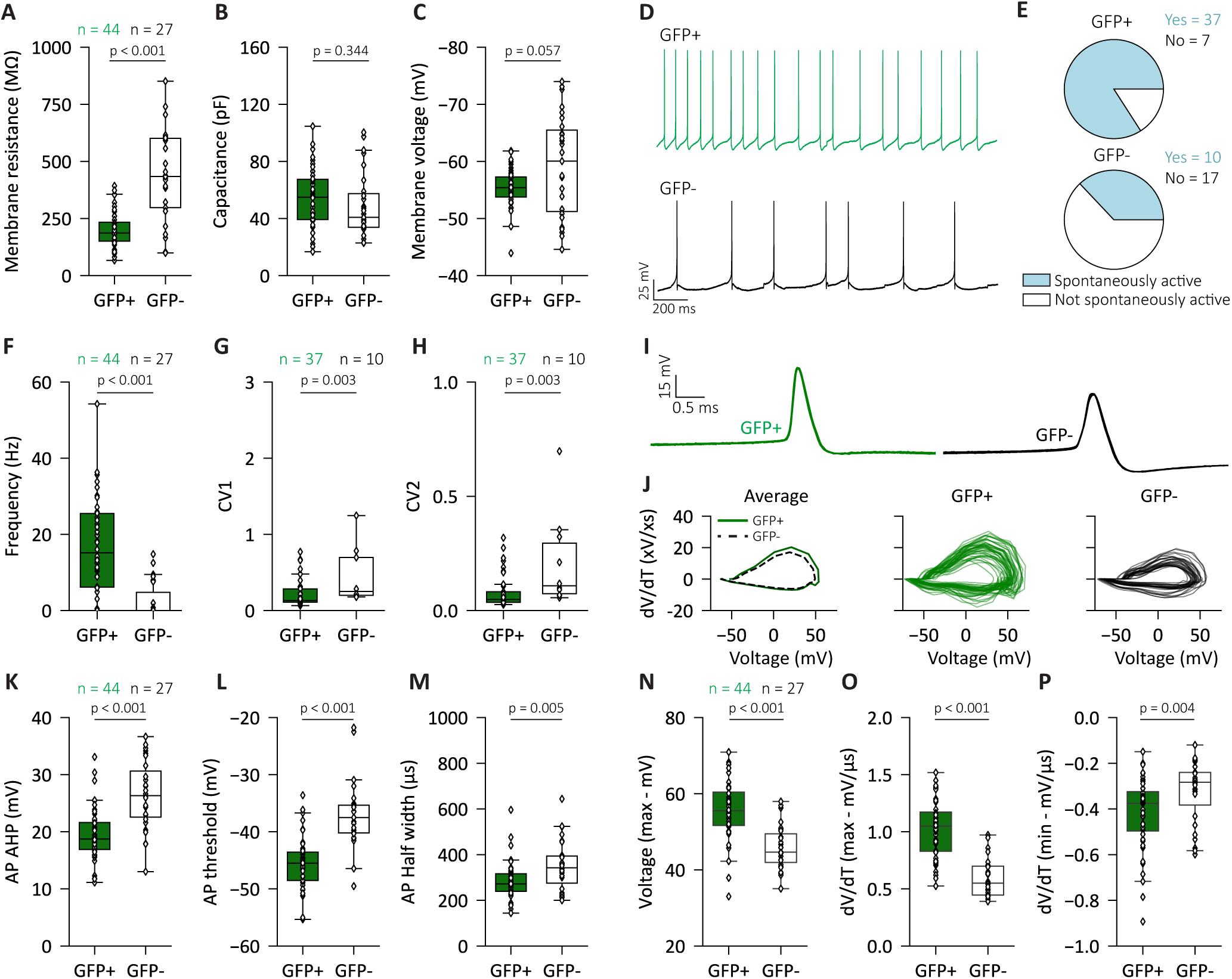
– Analyses of passive, spontaneous, and action potential properties of IO-projecting MDJ neurons. **(A-C)** Box plots of the membrane resistance (GFP+: 202.1 ± 78.2 MΩ, n = 44; GFP-: 439.7 ± 193.9 MΩ, n = 27; p = 1.34e-6, Welch’s t-test), membrane capacitance (GFP+: 54.0 ± 20.6 pF, n = 44; GFP-: 49.0 ± 22.5 pF, n = 27; p = 0.344, Independent t-test) and resting membrane voltage (GFP+: -55.4 ± 3.24 mV, n = 44; GFP-: -59.4 ± 8.67 mV, n = 27; p = 0.057, Welch’s t-test) of recorded GFP+ and GFP- neurons. **(D)** Example traces of spontaneous action potentials recorded in a GFP+ and GFP- neuron. **(E)** The proportions of GFP+ (37/44) and GFP- (10/27) neurons which are spontaneously active (p = 7.87e-5, Fisher’s exact test). **(F-H)** Boxplots of the spontaneous action potential frequency (GFP+: 15.2 (19.3) Hz, n = 44; GFP-: 0 (4.78) Hz, n = 27; p = 4.36e-7, Mann-Whitney U-test), global coefficient of variation (CV1) (GFP+: 0.130 (0.173), n = 37; GFP-: 0.250 (0.499), n = 10; p = 0.003, Mann-Whitney U-test), and local coefficient of variation (CV2) (CV2; GFP+: 0.048 (0.046), n = 37; GFP-: 0.108 (0.221), n = 10; p = 0.003, Mann-Whitney U-test). **(I)** Example traces of action potentials recorded in a GFP+ and GFP- neuron. A total of 10 action potentials recorded in the same neuron are overlayed. **(J)** Plots of the membrane voltage against the derivative of the membrane voltage (the change in voltage over time). Left shows the average for all GFP+ and GFP+ neurons. Middle and right show the distributions for GFP+ and GFP- neurons, respectively. **(K-M)** Boxplots of the spike afterhyperpolarization (GFP+: 18.7 (4.71) mV, n = 44 ; GFP-: 26.3 (8.06) mV, n = 27 ; p = 4.07e-6, Mann-Whitney U-test), action potential threshold (GFP+: -45.5 (4.94) mV, n = 44 ; GFP-: -37.1 (4.86) mV, n = 27 ; p = 7.81e-8, Mann-Whitney U-test) and action potential half width (GFP+: 272.0 (76.5) µs, n = 44 ; GFP-: 342.0 (119.0) µs, n = 27 ; p = 0.005, Mann-Whitney U-test). **(N-P)** Boxplots of the maximum voltage (GFP+: 55.8 ± 7.87 mV, n = 44; GFP-: 45.9 ± 5.68 mV, n = 27; p = 4.02e-7, Independent t-test), maximum voltage increase (Max dV/dT; GFP+: 1.02 ± 0.25 mV/µs, n = 44; GFP-: 0.59 ± 0.16 mV/µs, n = 27; p = 1.34e-12, Welch’s t-test) and maximum voltage decrease (Min dV/dT; GFP+:-0.38 ± 0.17 mV/µs, n = 44; GFP-:-0.28 ± 0.14 mV/µs, n = 27; p = 0.004, Welch’s t-test). Data are represented as mean ± SD.

Under current-clamp conditions, we observed that 37 out of 44 (84%) recorded GFP+ neurons fired action potentials spontaneously, in contrast to only 10 out 27 (37%) of the recorded GFP- neurons (p = 7.83e-5, Odds ratio = 8.99, Fisher’s exact test) (Fig 2d,e). GFP+ neurons fired more frequently (GFP+: 15.2 (19.3) Hz, n = 44; GFP-: 0 (4.78) Hz, n = 27; p = 4.36e-7, Mann-Whitney U-test) and with a lower coefficient of variation than GFP- neurons (GFP+: 0.130 (0.173), n = 37; GFP-: 0.250 (0.499), n = 10; p = 0.003, Mann-Whitney U-test) (Fig 2f,g). Similar results were obtained when the coefficient of variation was calculated for individual pairs of spikes (CV2; GFP+: 0.048 (0.046), n = 37; GFP-: 0.108 (0.221), n = 10; p = 0.003, Mann-Whitney U-test) (Fig 2g). Thus, GFP+ neurons showed a higher spontaneous spike frequency with lower inter-spike interval variability.

We also applied a slow ramping current to isolate individual action potentials for analysis of the active properties of GFP+ and GFP- neurons (Fig 2i,j). The action potentials of GFP+ neurons had smaller afterhyperpolarizations (GFP+: 18.7 (4.71) mV, n = 44 ; GFP-: 26.3 (8.06) mV, n = 27 ; p = 4.07e-6, Mann-Whitney U-test), lower spike thresholds (GFP+:-45.5 (4.94) mV, n = 44 ; GFP-:-37.1 (4.86) mV, n = 27 ; p = 7.81e-8, Mann-Whitney U-test), and shorter action potential half widths (GFP+: 272.0 (76.5) µs, n = 44 ; GFP-: 342.0 (119.0) µs, n = 27 ; p = 0.005, Mann-Whitney U-test) (Fig 2k-m). GFP+ neurons also on average reached a higher maximum membrane voltage on average (GFP+: 55.8 ± 7.87 mV, n = 44; GFP-: 45.9 ± 5.68 mV, n = 27; p = 4.02e-7, Independent t-test) (Fig 2n). Finally, the maximum and minimum derivatives of the membrane voltage during an action potential were larger in GFP+ neurons (Max dV/dT; GFP+: 1.02 ± 0.25 mV/µs, n = 44; GFP-: 0.59 ± 0.16 mV/µs, n = 27; p = 1.34e-12, Welch’s t-test) (Min dV/dT; GFP+:-0.38 ± 0.17 mV/µs, n = 44; GFP-:-0.28 ± 0.14 mV/µs, n = 27; p = 0.004, Welch’s t-test) (Fig 2o,p). Altogether, these results suggest that there are considerable differences in the active properties between IO- and non-IO projecting neurons, i.e., action potential dynamics were faster, both during the depolarization- and the re-polarization phase, and action potential threshold was lower for IO-projecting neurons.

Next, we investigated the action potential generation of IO-projecting (GFP+) and non-IO-projecting (GFP-) neurons during depolarizing current injections in current-clamp mode (Fig 3a). Neurons typically showed a higher initial action potential frequency and amplitude, followed by sustained firing at a lower frequency and amplitude (Fig 3b-d). Notably, the frequency of action potentials decreased more sharply for GFP- neurons over the course of depolarization (Time*GFP; F(1,59.305) = 8.1882, p = 0.0058, linear mixed model) (Fig 3b). We also observed a more pronounced decrease in action potential amplitude in GFP neurons over the course of depolarization (Time*GFP; F(1,43.074) = 7.1362, p = 0.0106, linear mixed model). In addition, we observed a higher average action potential rate in GFP+ neurons (F(1,71.720) = 4.4294, p = 0.0388, linear mixed model), as well as a shorter latency until the first action potential (F(1,41.776) = 10.003, p=0.0029, linear mixed model) (Fig 3d,e). After cessation of depolarizing current injections, both groups of neurons displayed an after-hyperpolarization, which had a larger amplitude in the IO-projecting GFP+ neurons (F(1,69.055) = 55.629, p = 1.943e-10, linear mixed model) (Fig 3f). Thus, IO-projecting neurons of the MDJ fired higher and faster to depolarizations and had larger after hyperpolarization than presumable non-IO-projecting neurons.

**Figure 3.**
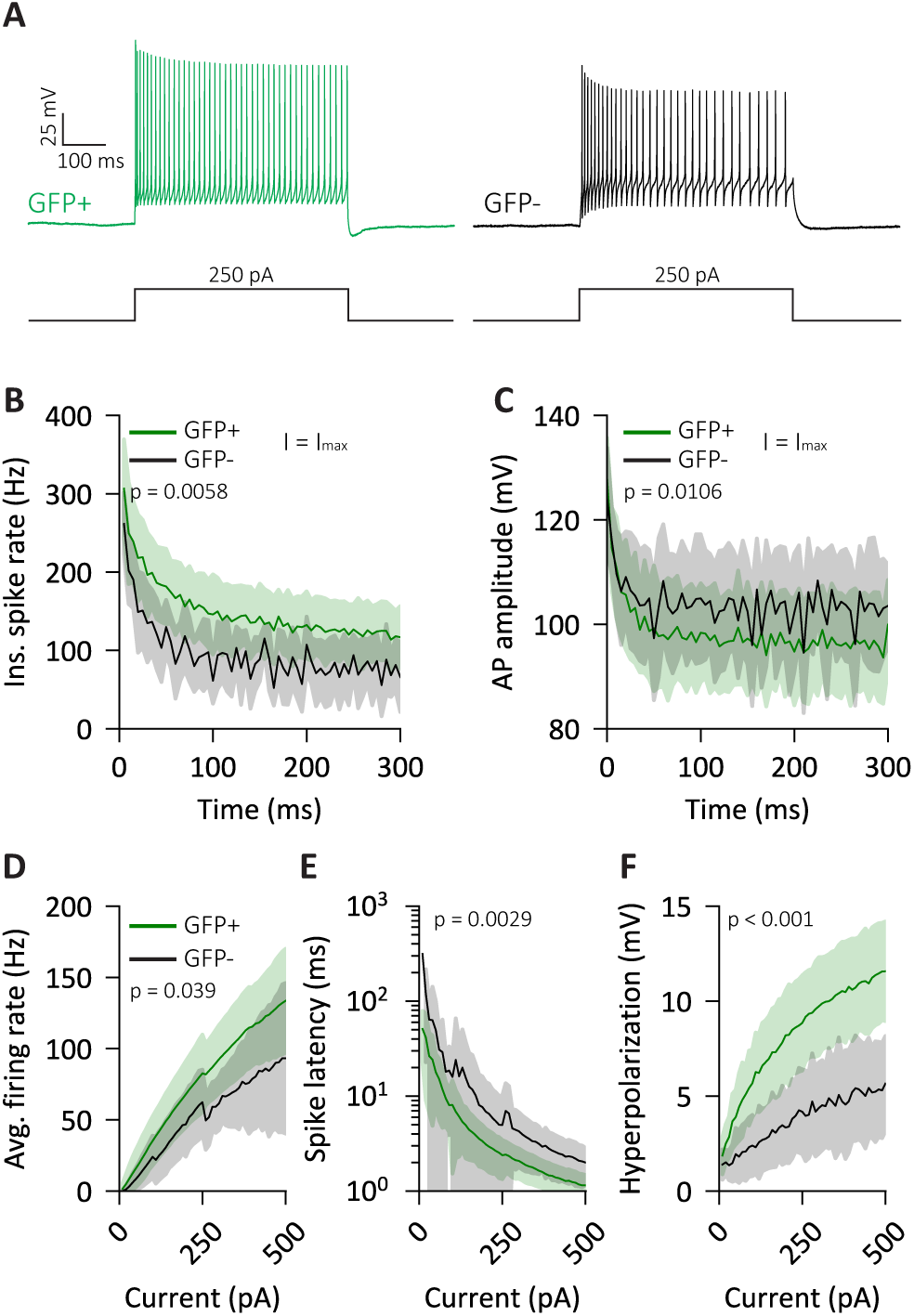
Response of IO- and non-IO projecting MDJ neurons to depolarizing currents. **(A)** Example traces of depolarizing current injections in GFP+ and GFP- neurons. **(B)** The instantaneous firing rate of GFP+ and GFP- neurons over time in response to a depolarizing current injection (Time*GFP; F(1,59.305) = 8.1882, p = 0.0058, linear mixed model). The current amplitude was the maximum current for each recorded neuron. **(C)** The attenuation of action potential (AP) amplitude over time in response to a depolarizing current injection (Time*GFP; F(1,43.074) = 7.1362, p = 0.0106, linear mixed model). The current amplitude was the maximum current for each recorded neuron. **(D-F)** Firing rate (F(1,71.720) = 4.4294, p = 0.0388, linear mixed model), spike latency (F(1,41.776) = 10.003, p=0.0029, linear mixed model) and hyperpolarization (F(1,69.055) = 55.629, p = 1.943e-10, linear mixed model) of GFP+ and GFP- neurons in response to increasingly large depolarizing current injections. The ‘skip’ in data at 250 pA is due to unequal group size (see methods). Data are represented as mean ± SD.

We next investigated the responses to hyperpolarizing current injections (Fig 4a). Almost all GFP+ neurons (43 out of 44 (98%) recorded GFP+ neurons) produced rebound action potentials upon cessations of hyperpolarizing current injections, whereas rebound action potentials could be induced in only 7 out of 27 (26%) recorded GFP- neurons (Fig 4b,c) (p = 7.19e-11, Fisher’s exact test). The number of rebound action potentials increases with the amplitude of the hyperpolarizing current, but this effect typically saturated at injected currents of around -250 pA (Fig 4d, all neurons). GFP+ neurons also fired significantly more rebound action potentials than GFP- neurons (F(1, 73.582) = 40.897, p = 1.314e-8, linear mixed model) (Fig 4d). Unlike depolarizing current injections, we observed no difference in the spike latency of rebound action potentials (F(1,49.509) = 2.7847, p = 0.1015, linear mixed model) (Fig 4e). Finally, hyperpolarizing currents induced an initial sag (Fig 4a, arrow) in the membrane voltage, which was more pronounced in GFP+ neurons (F(1,72.688) = 46.043, p = 2.640e-9, linear mixed model) (Fig 4f).

**Figure 4.**
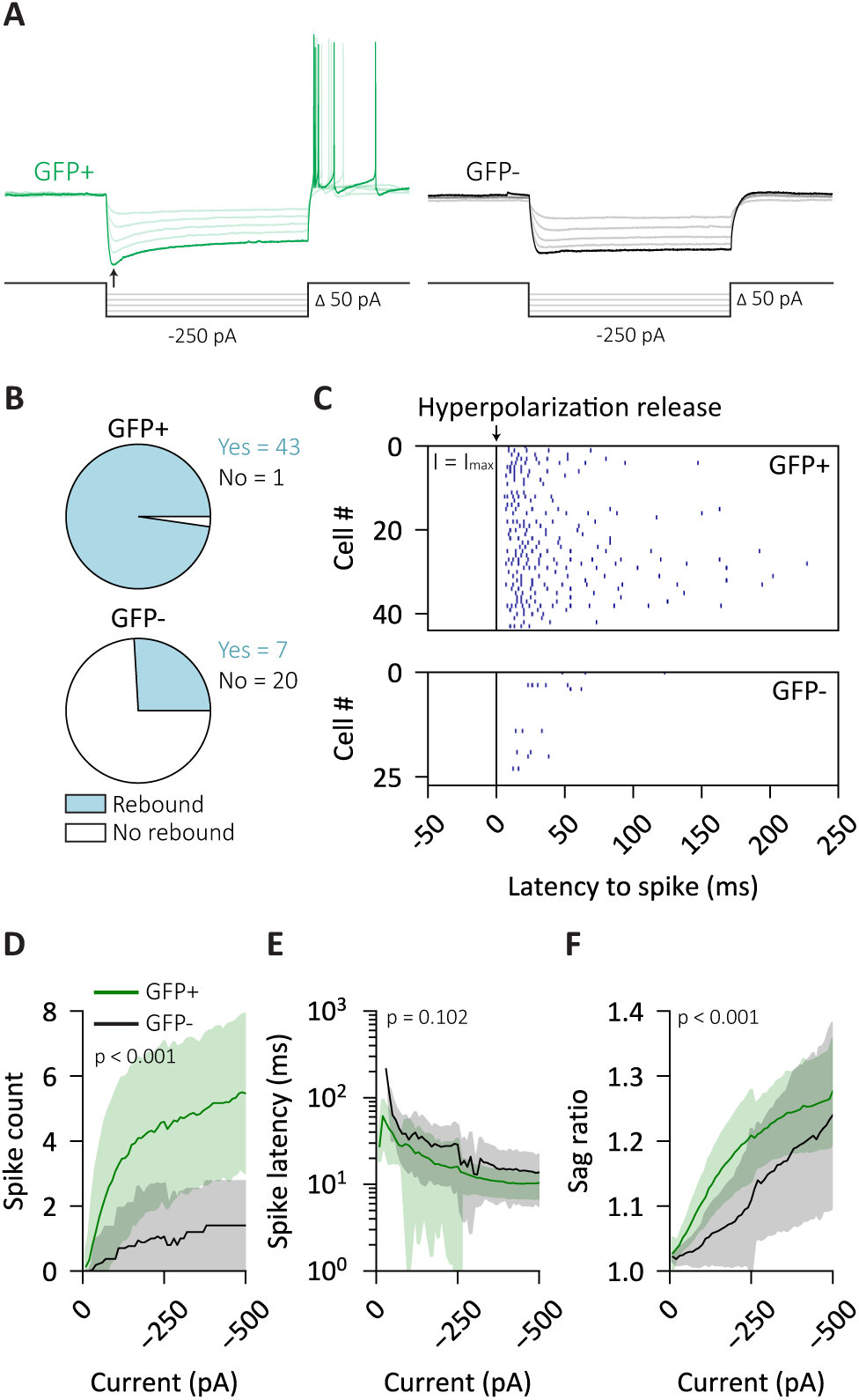
Response of IO- and non-IO projecting MDJ neurons to hyperpolarizing currents. **(A)** Example traces of hyperpolarizing current injections in GFP+ and GFP- neurons. **(B)** The proportions of GFP+ (43/44) and GFP- (7/27) neurons which fire rebound action potentials after the release of hyperpolarization (p = 7.19e-11, Fisher’s exact test). **(C)** Raster plot showing the timing of rAP firing after the release of hyperpolarization (t = 0) for all included GFP+ and GFP- neurons. The current amplitude was the maximum hyperpolarizing current for each recorded neuron. **(D-F)** Rebound spike count (F(1, 73.582) = 40.897, p = 1.314e-8, linear mixed model), spike latency (F(1,49.509) = 2.7847, p = 0.1015, linear mixed model) and sag ratio (F(1,72.688) = 46.043, p = 2.640e-9, linear mixed model) of GFP+ and GFP- neurons in response to increasingly large hyperpolarizing current injections. The ‘skip’ in data at 250 pA is due to unequal group size (see methods). Data are represented as mean ± SD.

### MDJ-IO neurons represent a physiologically distinct population

To further classify the physiological heterogeneity of MDJ-IO neurons, we performed principal component analysis based on their electrophysiological properties. GFP+ and GFP- neurons formed separated clusters in the PC1/PC2 parameter space (PC1 + PC2 variance explained = 65.9%, Fig 5a,b). K-means clustering effectively identified GFP+ and GFP- neurons as separate clusters, although 2 GFP- neurons were classified in cluster 1 and 2 GFP+ neuron in cluster 2 (Fig. 5a). This may suggest that these former five neurons do project to the IO, but did not express the retrograde virus, while the latter cell may have picked up off-target virus expression outside the IO.

**Figure 5.**
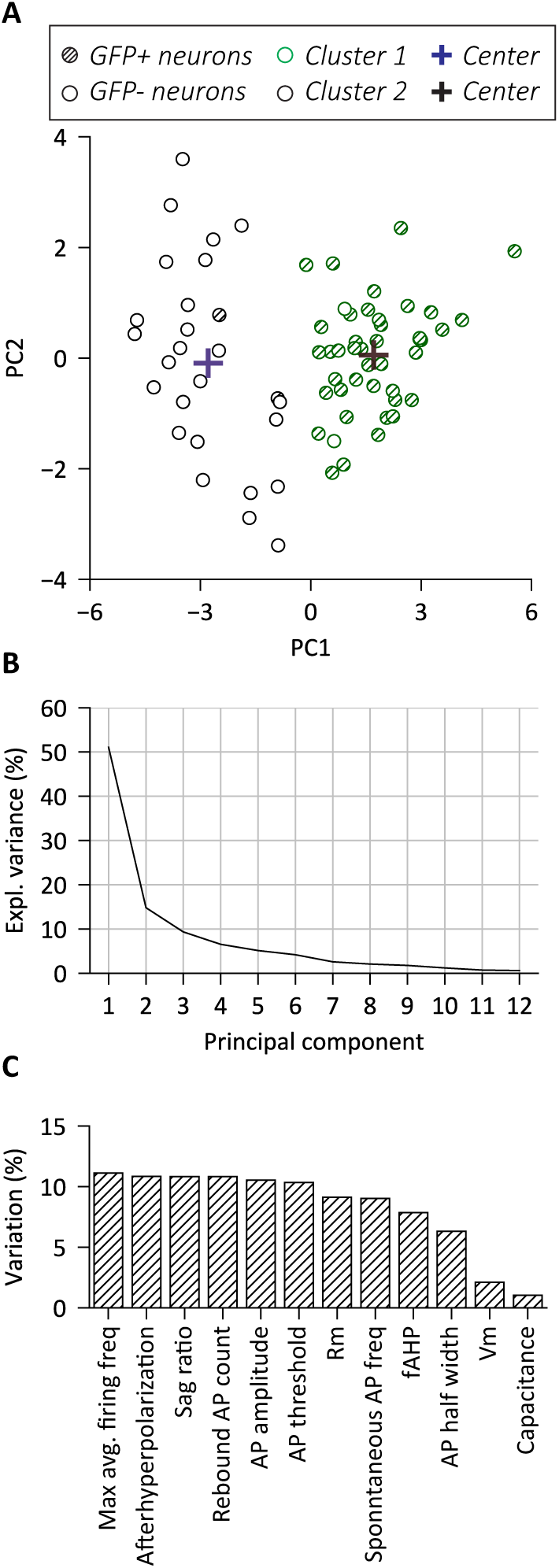
Principal component analysis and clustering of MDJ neuron electrophysiological properties. **(A)** 2D plot of the first two principal components, overlayed with the clusters identified by k-means clustering. **(B)** The explained variance of each principal component. **(C)** The variance explained by each of the original features put into the analysis in percentages.

Analysis of the eigenvalues that represent the contribution of each variable revealed that the differences between the two clusters did depend roughly equally on all variables, except membrane voltage and membrane capacitance (Fig 5c). In short, GFP+ neurons displayed an electrophysiological profile that can be distinguished from that of GFP- neurons.

### IPSC recordings in the MDJ

One of the most consistent and striking features we observed in the IO-projecting neurons was the firing of rebound action potentials, which is a feature typical for neurons receiving strong inhibitory inputs (Grenier et al., 1998; Aizenman & Linden, 1999; Hoebeek et al., 2010). This would be consistent with the inhibitory drive from, in particular, the entopeduncular nucleus to the MDJ (Ruigrok et al., 2023). Therefore, we quantified the inhibitory input to IO-projecting neurons. We recorded spontaneous, inhibitory postsynaptic currents (sIPSCs) from GFP+ neurons in the presence of blockers of AMPA (NBQX) and NMDA receptors (D(L)-AP5). For these recordings, we used an internal solution based on caesium chloride to invert the electrochemical gradient for chloride at a Vm of-65 mV.

All recorded GFP+ neurons showed sIPSCs (Fig 6–a,b) (frequency: 4.73 ± 2.09 Hz; amplitude: -62.5 ± 25.6 pA; rise time; 2.00 ± 0.31 ms; decay time: 5.27 ± 1.34 ms; n = 15 neurons from 5 mice). To confirm that these sIPSCs were GABAergic, we washed in the GABAA-receptor antagonist picrotoxin, which almost entirely abolished sIPSCs within 20 minutes (frequency; F(1,6.0687) = 156.47, p = 1.466e-5, linear mixed model) (Fig 6c,d). In addition, the average amplitude and rise time of sIPSCs was significantly reduced after the addition of picrotoxin (amplitude; F(1, 2.702) = 58.028, p = 0.0067; rise time; F(1, 4.9424) = 11.812, p = 0.0188, linear mixed model) (Fig 6d). No significant change was observed in the decay time in the presence of picrotoxin (F(1,4.9115) = 0.98111, p = 0.3682, linear mixed model). Consistent with these data, we also observed overlapping expression of VGAT and gephyrin around individual cell bodies around the fasciculus retroflexus (Fig 6e,f). These results indicate that IO-projecting neurons in the MDJ indeed receive GABAergic inhibitory input, potentially facilitating the firing of rebound action potentials.

**Figure 6.**
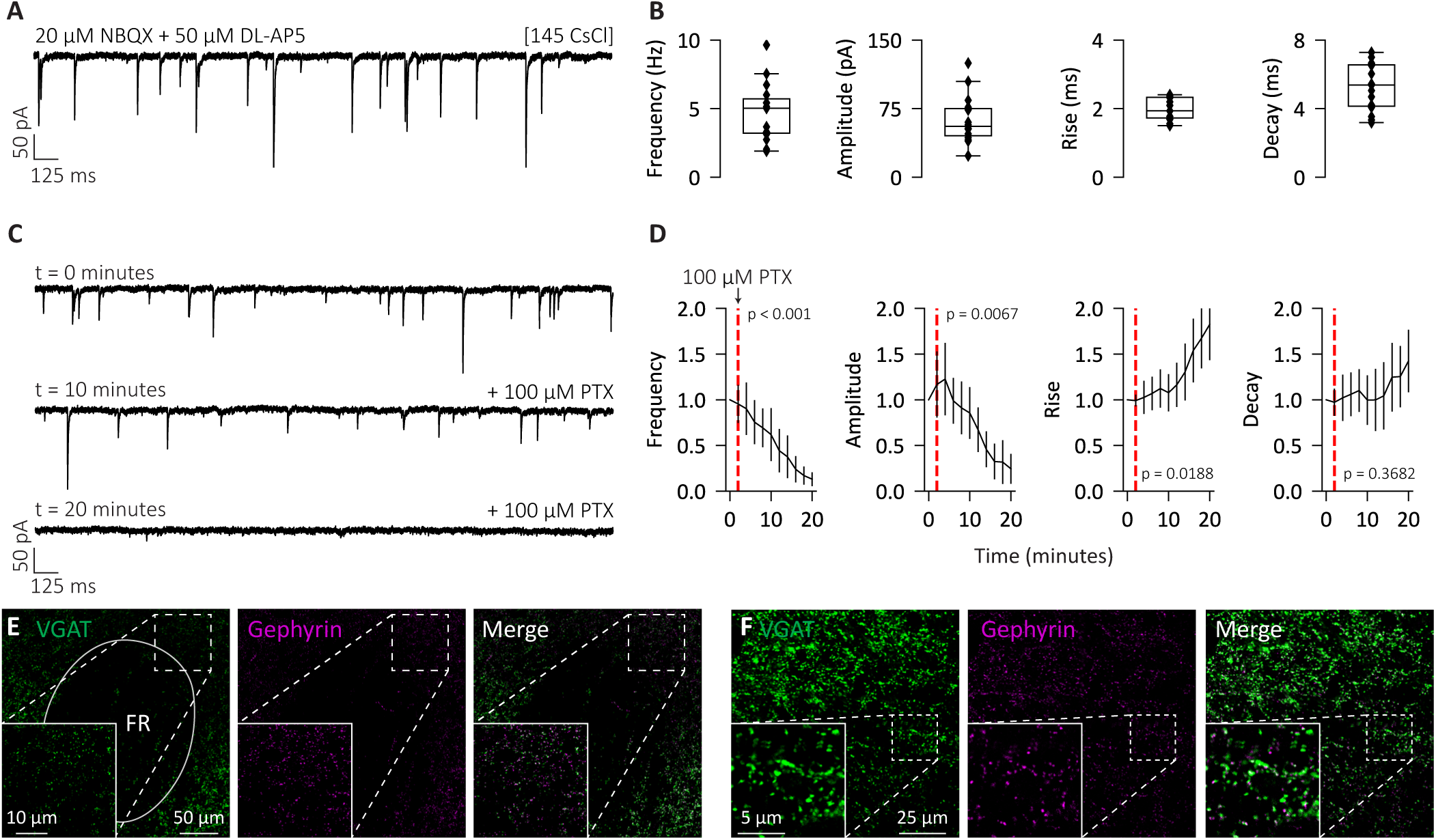
Inhibitory inputs to MDJ-IO neurons. **(A)** Example trace of sIPSC recordings performed with caesium chloride (CsCl) and in the presence of the AMPA and NMDA blockers NBQX and DL-AP5, respectively. **(B)** Boxplots of sIPSC frequency, amplitude, rise time, and decay time (frequency: 4.73 ± 2.09 Hz; amplitude:-62.5 ± 25.6 pA; rise time; 2.00 ± 0.31 ms; decay time: 5.27 ± 1.34 ms; n = 15 neurons from 5 mice). **(C)** Example traces of a recorded MDJ-IO neuron before (t=0), 10 minutes after (t=10), and 20 minutes (t=20) after the addition of the GABA receptor blocker picrotoxin (PTX). **(D)** Time-course of the frequency minutes (F(1,6.0687) = 156.47, p = 1.466e-5, linear mixed model), amplitude (F(1, 2.702) = 58.028, p = 0.0067, linear mixed model), rise time (F(1, 4.9424) = 11.812, p = 0.0188, linear mixed model) and decay time (F(1,4.9115) = 0.98111, p = 0.3682, linear mixed model) of sIPSCs after the addition of PTX (red dotted line). **(E-F)** Confocal images of the MDJ labelled with fluorescent markers for VGAT (left) and Gephyrin (middle), and overlayed together (right). The image in (e) is the same animal as in (f), but (f) is at a higher magnification and resolution. Data are represented as mean ± SD.

### Optogenetic stimulation of cortical and cerebellar inputs

The MDJ receives prominent input from the cerebral cortex and the cerebellar nuclei (Kubo et al., 2018; Wang et al., 2022; Ruigrok et al., 2023). Previous work has shown at the light microscopic level that terminals from the cerebellar nuclei and neocortical areas converge onto individual IO-projecting neurons of the MDJ (Wang et al., 2022). However, to date it is unclear whether both inputs are actually functional at the level of individual MDJ neurons, and it is unclear to what extent both types of input are excitatory or inhibitory and how they interact. To test the ability of the cerebral cortex and the cerebellar nuclei to evoke postsynaptic responses in MDJ neurons, we made viral injections to express the opsins ChrimsonR and Chronos in the cerebral cortex and cerebellar nuclei, respectively (Fig 7a,b). Given the widespread distribution of IO-projecting neurons in the cerebral cortex, we made multiple injections, targeting both motor (M1 and M2) and somatosensory areas (S1 and S2). Injections in the DCN were primarily targeting the dentate and interposed nuclei. To identify the IO-projecting neurons, we made the same retrograde AAV injections in the IO as described above.

**Figure 7.**
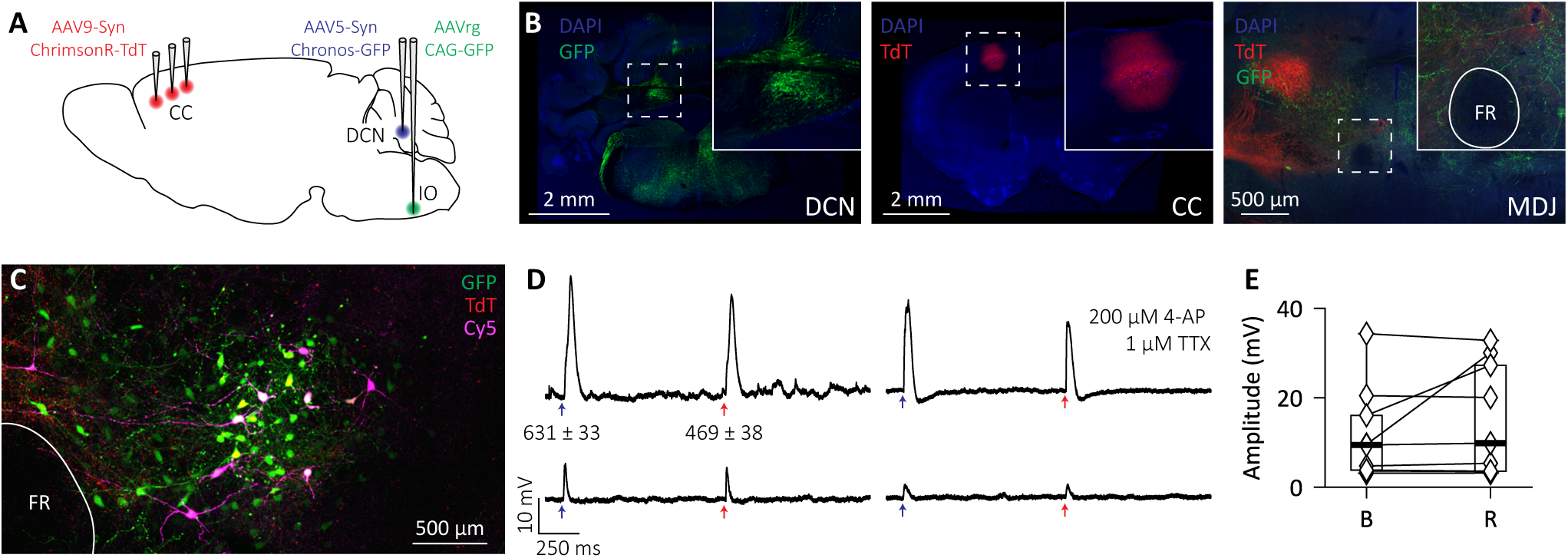
Optogenetic stimulation of DCN and CC afferents to the MDJ. **(A)** Viral injections used in the optogenetics experiments. AAV9-Syn-ChrimsonR-TdT was injected in the CC, specifically M1/M2 and S1/S2. AAV5-Syn-Chronos-GFP was injected in the DCN. AAVrg-CAG-GFP was injected in the IO. **(B)** Confocal images with examples of viral expression in the DCN (left), CC (middle), and MDJ (right). Note that the AAVrg injection in the IO was omitted for this example. **(C)** Confocal image of biocytin filled neurons recorded during optogenetic experiments.

At least three weeks following injections, we performed dual-optogenetic stimulation of DCN and cerebral cortical axons, while performing whole-cell recordings from GFP+ neurons in acute brain slices (Fig 7c). Recordings were made in the presence of TTX and 4-AP to isolate optogenetically activated synapses. We found responses to the activation of both opsins in 9/42 neurons recorded neurons, or in 21% (Fig 7d,e; Fig 8). All observed responses from the cerebellar nuclei and cerebral cortex were excitatory. Thus, we physiologically show that excitatory synaptic terminals from the cerebral cortex and the cerebellar nuclei converge on single MDJ-IO neurons, but it remains to be determined how the inhibition in the MDJ comes about.

**Figure 8.**
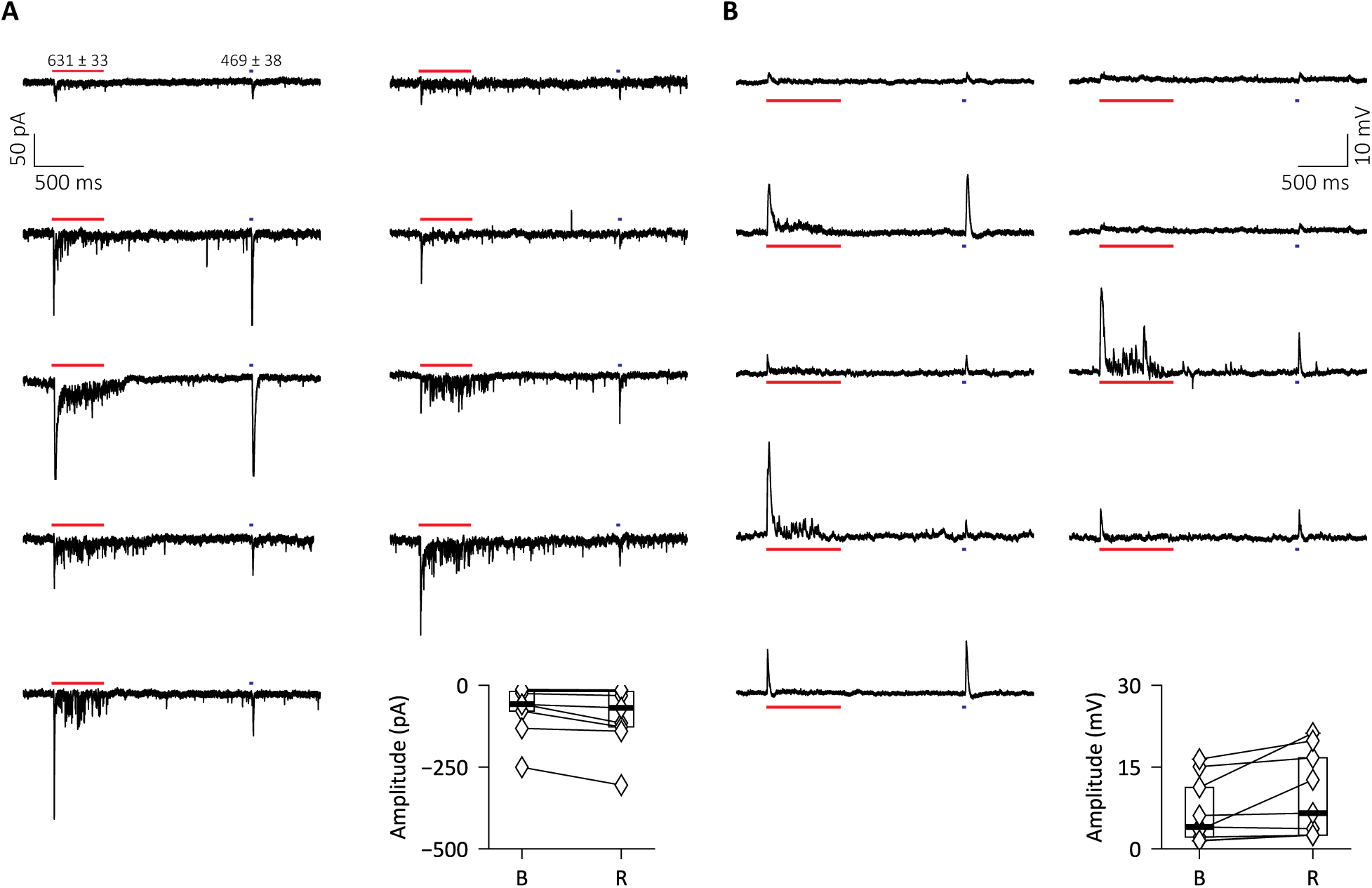
Optogenetic responses of all GFP+ neurons with dual responses after depletion of ChrimsonR fibres. **(A)** Dual optogenetic responses to 500 ms of red light followed by a brief blue light stimulation in voltage clamp. Plot shows the average amplitude of the response for each cell to blue (B) and red (R) light stimulation. **(B)** Same as in (A) but in current-clamp.

## Discussion

In this study, we characterised the electrophysiological properties of MDJ neurons that specifically project to the IO. We found that, in the MDJ, IO-projecting neurons were intermingled with non-IO-projecting neurons. IO-projecting MDJ neurons were distinct from their putative non-IO projecting neighbours and could be distinguished based on these electrophysiological properties. In particular, we found that IO-projecting neurons spontaneously fire action potentials, respond robustly to depolarizing currents, and fire rebound action potentials in response to hyperpolarizing currents. We also show that IO-projecting neurons receive inhibitory input, suggesting that they may be bidirectionally regulated by GABAergic inhibition and glutamatergic excitation. Excitatory synaptic input to the MDJ can come from the cerebellar nuclei and cerebral cortex, and we show that these inputs can converge at the level of individual neurons. Altogether, these results indicate that IO-projecting MDJ neurons consist of a physiologically distinct and homogeneous population, which integrates excitatory input from the cerebral cortex and the DCN, as well as inhibitory input from a currently unknown source. Given their intrinsic properties and combined external inputs, MDJ neurons are well suited to engage both temporal and rate coding mechanisms to control timing of the olivary neurons, a feature that may help the olivocerebellar system to optimize spike timing-dependent plasticity during learning.

### Experimental considerations

As the anatomical borders of the nuclei composing the MDJ are poorly defined in rodents, we relied on a viral tracing approach to identify IO-projecting MDJ neurons. To this end, we injected retrograde GFP-expressing virus bilaterally in the IO prior to the electrophysiological recordings. Due to the deep location and small size of the IO, we cannot rule out that some viral particles unintendedly ended up in adjacent regions. Conversely, not all MDJ fibres will have taken up the virus. Although we excluded mice in which the virus injections were visibly off-target, it is likely that our classification of IO-projecting and non-IO-projecting neurons based on the retrograde tracing was imperfect. However, the two clusters of neurons that were found in the principal component and clustering analysis matched well with the expression pattern of GFP, indicating that the identification of IO-projecting and non-IO-projecting neurons in the MDJ was generally reliable.

An important finding in our study was the presence of rebound action potentials in IO-projecting MDJ neurons. The ionic conductances that facilitate the firing of rebound action potentials in MDJ neurons were not identified in this study. Typically, rebound firing is induced by T-type Ca2+ channels and/or hyperpolarization activated cation channels triggering Ih currents (Aizenman & Linden, 1999; Surges et al., 2006). It seems likely that Ih currents play a significant role in the generation of rebound firing in IO-projecting MDJ neurons, as we observe an initial voltage sag when hyperpolarization is applied, typical of neurons with strong Ih currents (Robinson & Siegelbaum, 2003). A caveat of this conclusion is that, because we recorded rebound action potentials from a starting membrane voltage of-65 mV, Ih currents may be in a state of higher conductance. At resting membrane potentials, rebound firing may thus be less prominent than we observed. However, it is entirely possible that synchronised inhibition, mediated through the GABAergic inputs we identified in the MDJ, may also strongly hyperpolarize MDJ neurons in vivo, thus eliciting rebound spikes comparable to what we observe in vitro. Further experiments will have to validate the functional occurrence and relevance of rebound firing, as well as the influence of inhibition on eliciting these spikes during behaviour.

### Information coding in the MDJ

Even though the MDJ receives synaptic input from a large variety of brain regions (Onodera, 1984; Kubo et al., 2018; Wang et al., 2022; Ruigrok et al., 2023), we found that IO-projecting MDJ neurons are largely homogeneous in terms of their electrophysiological properties. Typically, IO-projecting MDJ neurons are spontaneously active. As such, the IO-projecting neurons in the MDJ can contribute to the suppression of intrinsic oscillatory activity and synchrony in the IO, as predicted by a biophysically plausible network model of the IO (Negrello et al., 2019). The tonic excitatory input from the MDJ may serve as contextual information, setting a baseline level of activity in the IO, while suppressing spontaneous oscillations and synchrony within the IO.

This latter role is in line with our observation that the I-F (current-frequency) curve in response to depolarizing inputs is mostly linear, which contrasts the flatter I-F curve of hyperpolarizing currents. The response to depolarizing current thus precisely transforms its input into a proportional output, suggesting that excitatory inputs to IO-projecting neurons can in principle be readily transformed into a rate code. In contrast, the rebound response is less dependent on input amplitude, yet more to the timed offset of inhibition, suggesting that the inhibitory inputs to IO-projecting MDJ neurons can also be transformed into a temporal code.

Furthermore, we also observe that depolarizing inputs generate brief high-frequency bursts of action potentials in IO-projecting MDJ neurons. High frequency burst activity in MDJ neurons may serve to increase the probability of firing in specific groups of IO neurons as well as setting the phase for a period of ∼200 ms (Negrello et al., 2019; Loyola et al., 2023). The exact phase of this oscillation will then also set up a window, in which subsequent excitatory signals will have a higher or lower probability of inducing spikes in the IO (Negrello et al., 2019).

The IO-projecting neurons of the MDJ receive substantial GABAergic inhibition. The origins of these inhibitory fibres are not fully known, but they are likely to include the entopeduncular nucleus that is an output nucleus of the basal ganglia (Ruigrok et al., 2023). The basal ganglia provide tonic inhibition at rest in order to suppress unwanted (motor) activity (Grillner & Robertson, 2016). At least in our in vitro preparation, GABAergic inhibition of IO-projecting MDJ neurons is not strong enough to prevent spiking at rest, it can still be assumed that inhibition from the basal ganglia can affect spiking activity in vivo. In particular, the end of inhibition could contribute to rebound spiking and thus to increased excitation of the IO.

Combining all these ideas, we propose the following theory. IO-projecting MDJ neurons fire regular action potentials at rest, but can rapidly and bidirectionally respond to depolarizing and hyperpolarizing inputs with high frequency bursts or spike-pauses followed by rebound firing, respectively. In vivo this could result in tonic regular activity is interrupted by high frequency bursts in response to a depolarizing input, for example from the cerebellar nuclei or cerebral cortex, or both simultaneously. The exact frequency of these bursts is highly dependent on the amplitude of the excitatory input, but also on the degree of inhibition preceding the excitation. Hyperpolarizing inputs may lead to a brief pause of spontaneous firing, followed by rebound firing, of which the timing and frequency is less dependent on stimulus intensity, but critically dependent on stimulus offset. Together, this allows IO-projecting MDJ neurons to generate both reliable and as well-timed signals based on its current activity level, direction of neurotransmission, and input intensity.

### Future directions and concluding remarks

Our physiological and histological evidence suggests that IO-projecting MDJ neurons receive GABAergic inhibitory input which, as we discussed, could be a physiological trigger for rebound action potentials analogous to the rebound spikes observed in cerebellar nuclei neurons in response to synchronized spike pauses of Purkinje cells (Witter et al., 2013). However, there are several remaining questions that we were unable to resolve in the present study. Additional identification of inhibitory pathways inside or to the MDJ, followed by physiological characterizations of these pathways will be needed to clarify the impact of GABAergic inhibition of IO-projection MDJ neurons. Furthermore, our study did not clarify the biophysical mechanisms that facilitate spontaneous activity, burst firing, rebound firing, or action potential dynamics. Future work should focus on further identifying the factors that determine the electrophysiological characteristics of these neurons. However, it is also of importance to move beyond ex vivo work and examine the firing dynamics of IO-projecting MDJ neurons and their effects on IO firing in vivo, preferably in awake behaving animals.

## Funding information

Financial support was provided by the Netherlands Organization for Scientific Research (NWO-ALW 824.02.001; CIDZ), the Dutch Organization for Medical Sciences (ZonMW 91120067; CIDZ), Medical Neuro-Delta (MD 01092019-31082023; CIDZ), INTENSE LSH-NWO (TTW/00798883; CIDZ), ERC-adv (GA-294775 CIDZ); The NIN Vriendenfonds for Albinism (CIDZ) as well as the Dutch NWO Gravitation Program, Dutch Brain Interface Initiative (DBI2 grant no. 024.005.022; CIDZ).

## Additional information

### Conflict of interests

The authors declare no conflicts of interest.

### Author contributions

The experiments were performed in the laboratory of Prof. Dr. Chris. I. De Zeeuw at the Erasmus Medical Centre, Rotterdam, The Netherlands. SV, XW, RB, LWJB, and CIDZ designed the study. SV and XW were involved in acquisition of the data. Analysis of data was performed by SV. SV, XW, RB, LWJB and CIDZ were involved in the interpretation of data. All authors were involved in drafting or revising the manuscript for critically important intellectual content. All authors have read and approved the final version of the manuscript and agree to be accountable for all aspects of the work in ensuring that questions related to the accuracy or integrity of any part of the work are appropriately investigated and resolved. All persons designated as authors qualify for authorship, and all those who qualify for authorship are listed.

## Acknowledgements

The authors thank Erika H. Sabel-Goedknegt for her assistance with the immunohistochemistry.

## Notes

### Competing Interest Statement

The authors have declared no competing interest.

## References

Aizenman CD & Linden DJ (1999). Regulation of the Rebound Depolarization and Spontaneous Firing Patterns of Deep Nuclear Neurons in Slices of Rat Cerebellum. Journal of Neurophysiology 82, 1697–1709.

Apps R & Watson TC (2013). Cerebro-Cerebellar Connections. In Handbook of the Cerebellum and Cerebellar Disorders, ed. Manto M, Schmahmann JD, Rossi F, Gruol DL & Koibuchi N, pp. 1131–1153. Springer Netherlands, Dordrecht. Available at: https://link.springer.com/10.1007/978-94-007-1333-8_48 [Accessed January 30, 2026].

Bazzigaluppi P, De Gruijl JR, Van Der Giessen RS, Khosrovani S, De Zeeuw CI & De Jeu MTG (2012). Olivary subthreshold oscillations and burst activity revisited. Front Neural Circuits; DOI: 10.3389/fncir.2012.00091.

Best AR & Regehr WG (2009). Inhibitory Regulation of Electrically Coupled Neurons in the Inferior Olive Is Mediated by Asynchronous Release of GABA. Neuron 62, 555–565.

Bouvier G, Aljadeff J, Clopath C, Bimbard C, Ranft J, Blot A, Nadal JP, Brunel N, Hakim V & Barbour B (2018). Cerebellar learning using perturbations. Elife.

Broersen R & De Zeeuw CI (2024). Keeping track of time: An interaction of mossy fibers and climbing fibers. Neuron 112, 2664–2666.

Choi S, Yu E, Kim D, Urbano FJ, Makarenko V, Shin H-S & Llinás RR (2010). Subthreshold membrane potential oscillations in inferior olive neurons are dynamically regulated by P/Q- and T-type calcium channels: a study in mutant mice: Membrane oscillations in mutant mice. The Journal of Physiology 588, 3031–3043.

Coesmans M, Weber JT, De Zeeuw CI & Hansel C (2004). Bidirectional parallel fiber plasticity in the cerebellum under climbing fiber control. Neuron 44, 691–700.

De Gruijl JR, Hoogland TM & De Zeeuw CI (2014). Behavioral Correlates of Complex Spike Synchrony in Cerebellar Microzones. Journal of Neuroscience 34, 8937–8947.

De Zeeuw CI, Hoebeek FE, Bosman LWJ, Schonewille M, Witter L & Koekkoek SK (2011). Spatiotemporal firing patterns in the cerebellum. Nat Rev Neurosci 12, 327–344.

De Zeeuw CI, Holstege JC, Ruigrok TJH & Voogd J (1989). Ultrastructural study of the GABAergic, cerebellar, and mesodiencephalic innervation of the cat medial accessory olive: Anterograde tracing combined with immunocytochemistry. J of Comparative Neurology 284, 12–35.

De Zeeuw CI, Hoogenraad CC, Koekkoek SKE, Ruigrok TJH, Galjart N & Simpson JI (1998). Microcircuitry and function of the inferior olive. Trends in Neurosciences 21, 391–400.

De Zeeuw CI & Ten Brinke MM (2015). Motor Learning and the Cerebellum. Cold Spring Harb Perspect Biol 7, a021683.

Eccles JC, Llinás R & Sasaki K (1966). The excitatory synaptic action of climbing fibres on the Purkinje cells of the cerebellum. The Journal of Physiology 182, 268–296.

Fredette BJ & Mugnaini E (1991). The GABAergic cerebello-olivary projection in the rat. Anat Embryol 184, 225–243.

Gao Z, Van Beugen BJ & De Zeeuw CI (2012). Distributed synergistic plasticity and cerebellar learning. Nat Rev Neurosci 13, 619–635.

Grenier F, Timofeev I & Steriade M (1998). Leading role of thalamic over cortical neurons during postinhibitory rebound excitation. Proc Natl Acad Sci USA 95, 13929–13934.

Grillner S & Robertson B (2016). The Basal Ganglia Over 500 Million Years. Current Biology 26, R1088–R1100.

Guo D & Uusisaari MY (2025). In vivo imaging of inferior olive neurons reveals roles of co-activation and cerebellar feedback in olivocerebellar signaling. ; DOI: 10.7554/eLife.105305.1. Available at: https://elifesciences.org/reviewed-preprints/105305v1 [Accessed January 23, 2026].

Hoebeek FE, Witter L, Ruigrok TJH & De Zeeuw CI (2010). Differential olivo-cerebellar cortical control of rebound activity in the cerebellar nuclei. Proc Natl Acad Sci USA 107, 8410–8415.

Hull C (2020). Prediction signals in the cerebellum: Beyond supervised motor learning. eLife 9, e54073.

Ikezoe K, Hidaka N, Manita S, Murakami M, Tsutsumi S, Isomura Y, Kano M & Kitamura K (2023). Cerebellar climbing fibers multiplex movement and reward signals during a voluntary movement task in mice. Commun Biol 6, 924.

Ito M, Sakurai M & Tongroach P (1982). Climbing fibre induced depression of both mossy fibre responsiveness and glutamate sensitivity of cerebellar Purkinje cells. The Journal of Physiology 324, 113–134.

Keser Z, Hasan KM, Mwangi BI, Kamali A, Ucisik-Keser FE, Riascos RF, Yozbatiran N, Francisco GE & Narayana PA (2015). Diffusion tensor imaging of the human cerebellar pathways and their interplay with cerebral macrostructure. Front Neuroanat; DOI: 10.3389/fnana.2015.00041.

Khosrovani S, Van Der Giessen RS, De Zeeuw CI & De Jeu MTG (2007). In vivo mouse inferior olive neurons exhibit heterogeneous subthreshold oscillations and spiking patterns. Proc Natl Acad Sci USA 104, 15911–15916.

Klapoetke NC et al. (2014). Independent optical excitation of distinct neural populations. Nat Methods 11, 338–346.

Kubo R, Aiba A & Hashimoto K (2018). The anatomical pathway from the mesodiencephalic junction to the inferior olive relays perioral sensory signals to the cerebellum in the mouse. The Journal of Physiology 596, 3775–3791.

Lang EJ (2002). GABAergic and Glutamatergic Modulation of Spontaneous and Motor-Cortex-Evoked Complex Spike Activity. Journal of Neurophysiology 87, 1993–2008.

Lang EJ, Sugihara I & Llinas R (1996). GABAergic modulation of complex spike activity by the cerebellar nucleoolivary pathway in rat. Journal of Neurophysiology 76, 255–275.

Llinas R, Baker R & Sotelo C (1974). Electrotonic coupling between neurons in cat inferior olive. Journal of Neurophysiology 37, 560–571.

Loyola S, Bosman LWJ, De Gruijl JR, De Jeu MTG, Negrello M, Hoogland TM & De Zeeuw CI (2019). Inferior Olive: All Ins and Outs. In Handbook of the Cerebellum and Cerebellar Disorders, ed. Manto M, Gruol D, Schmahmann J, Koibuchi N & Sillitoe R, pp. 1–56. Springer International Publishing, Cham. Available at: http://link.springer.com/10.1007/978-3-319-97911-3_43-2 [Accessed January 29, 2026].

Loyola S, Hoogland TM, Hoedemaker H, Romano V, Negrello M & De Zeeuw CI (2023). How inhibitory and excitatory inputs gate output of the inferior olive. eLife 12, e83239.

Negrello M, Warnaar P, Romano V, Owens CB, Lindeman S, Iavarone E, Spanke JK, Bosman LWJ & De Zeeuw CI (2019). Quasiperiodic rhythms of the inferior olive ed. Battaglia FP. PLoS Comput Biol 15, e1006475.

Onodera S (1984). Olivary projections from the mesodiencephalic structures in the cat studied by means of axonal transport of horseradish peroxidase and tritiated amino acids. J of Comparative Neurology 227, 37–49.

Palesi F, De Rinaldis A, Castellazzi G, Calamante F, Muhlert N, Chard D, Tournier JD, Magenes G, D’Angelo E & Gandini Wheeler-Kingshott CAM (2017). Contralateral cortico-ponto-cerebellar pathways reconstruction in humans in vivo: implications for reciprocal cerebro-cerebellar structural connectivity in motor and non-motor areas. Sci Rep 7, 12841.

Pernía-Andrade AJ, Goswami SP, Stickler Y, Fröbe U, Schlögl A & Jonas P (2012). A deconvolution-based method with high sensitivity and temporal resolution for detection of spontaneous synaptic currents in vitro and in vivo. Biophys J 103, 1429–1439.

Raymond JL, Lisberger SG & Mauk MD (1996). The Cerebellum: A Neuronal Learning Machine? Science 272, 1126–1131.

Robinson RB & Siegelbaum SA (2003). Hyperpolarization-Activated Cation Currents: From Molecules to Physiological Function. Annu Rev Physiol 65, 453–480.

Romano V, De Propris L, Bosman LW, Warnaar P, Ten Brinke MM, Lindeman S, Ju C, Velauthapillai A, Spanke JK, Midden-dorp Guerra E, Hoogland TM, Negrello M, D’Angelo E & De Zeeuw CI (2018). Potentiation of cerebellar Purkinje cells facilitates whisker reflex adaptation through increased simple spike activity. eLife 7, e38852.

Ruigrok TJH, Sillitoe RV & Voogd J (2015). Cerebellum and Cerebellar Connections. In The Rat Nervous System, pp. 133–205. Elsevier. Available at: https://linkinghub.elsevier.com/retrieve/pii/B9780123742452000097 [Accessed January 29, 2026].

Ruigrok TJH, Wang X, Sabel-Goedknegt E, Coulon P & Gao Z (2023). A disynaptic basal ganglia connection to the inferior olive: potential for basal ganglia influence on cerebellar learning. Front Syst Neurosci 17, 1176126.

Sasaki K, Oka H, Matsuda Y, Shimono T & Mizuno N (1975). Electrophysiological studies of the projections from the parietal association area to the cerebellar cortex. Exp Brain Res; DOI: 10.1007/BF00238732.

Sotelo C, Llinas R & Baker R (1974). Structural study of inferior olivary nucleus of the cat: morphological correlates of electrotonic coupling. Journal of Neurophysiology 37, 541–559.

Surges R, Sarvari M, Steffens M & Els T (2006). Characterization of rebound depolarization in hippocampal neurons. Biochemical and Biophysical Research Communications 348, 1343–1349.

Suvrathan A, Payne HL & Raymond JL (2016). Timing Rules for Synaptic Plasticity Matched to Behavioral Function. Neuron 92, 959–967.

Suzuki L, Coulon P, Sabel-Goedknegt EH & Ruigrok TJH (2012). Organization of Cerebral Projections to Identified Cerebellar Zones in the Posterior Cerebellum of the Rat. J Neurosci 32, 10854–10869.

Swenson RS, Sievert CF, Terreberry RR, Neafsey EJ & Castro AJ (1989). Organization of cerebral cortico-olivary projections in the rat. Neuroscience Research 7, 43–54.

Timmann D, Drepper J, Frings M, Maschke M, Richter S, Gerwig M & Kolb FP (2010). The human cerebellum contributes to motor, emotional and cognitive associative learning. A review. Cortex 46, 845–857.

Tsutsumi S, Hidaka N, Isomura Y, Matsuzaki M, Sakimura K, Kano M & Kitamura K (2019). Modular organization of cerebellar climbing fiber inputs during goal-directed behavior. eLife 8, e47021.

Van Der Giessen RS, Koekkoek SK, Van Dorp S, De Gruijl JR, Cupido A, Khosrovani S, Dortland B, Wellershaus K, Degen J, Deuchars J, Fuchs EC, Monyer H, Willecke K, De Jeu MTG & De Zeeuw CI (2008). Role of Olivary Electrical Coupling in Cerebellar Motor Learning. Neuron 58, 599–612.

Vrieler N, Loyola S, Yarden-Rabinowitz Y, Hoogendorp J, Medvedev N, Hoogland TM, De Zeeuw CI, De Schutter E, Yarom Y, Negrello M, Torben-Nielsen B & Uusisaari MY (2019). Variability and directionality of inferior olive neuron dendrites revealed by detailed 3D characterization of an extensive morphological library. Brain Struct Funct 224, 1677–1695.

Wang X, Novello M, Gao Z, Ruigrok TJH & De Zeeuw CI (2022). Input and output organization of the mesodiencephalic junction for cerebro-cerebellar communication. J of Neuroscience Research 100, 620–637.

Witter L, Canto CB, Hoogland TM, De Gruijl JR & De Zeeuw CI (2013). Strength and timing of motor responses mediated by rebound firing in the cerebellar nuclei after Purkinje cell activation. Front Neural Circuits; DOI: 10.3389/fncir.2013.00133.

